# Continuum Approximation of Invasion Probabilities

**DOI:** 10.1101/213348

**Authors:** Rebecca K. Borchering, Scott A. McKinley

**Affiliations:** Department of Biology; Emerging Pathogens Institute; and Department of Mathematics, University of Florida, Gainesville, FL.; Department of Mathematics, Tulane University, New Orleans, LA.

**Keywords:** invasion probability, stochastic population model, diffusion approximation

## Abstract

In the last decade there has been growing criticism of the use of Stochastic Differential Equations (SDEs) to approximate discrete state-space, continuous-time Markov chain population models. In particular, several authors have demonstrated the failure of *Diffusion Approximation*, as it is often called, to approximate expected extinction times for populations that start in a quasi-stationary state.

In this work we investigate a related, but distinct, population dynamics property for which Diffusion Approximation fails: invasion probabilities. We consider the situation in which a few individual are introduced into a population and ask whether their collective lineage can successfully invade. Because the population count is so small during the critical period of success or failure, the process is intrinsically stochastic and discrete. In addition to demonstrating how and why the Diffusion Approximation fails in the large population limit, we contrast this analysis with that of a sometimes more successful alternative WKB-like approach. Through numerical investigations, we also study how these approximations perform in an important intermediate regime. In a surprise, we find that there are times when the Diffusion Approximation performs well: particularly when parameters are near-critical and the population size is small to intermediate.

## 1. Introduction

Invasion events are fundamental in population biology. In the study of disease there are interesting examples both at the cellular and whole-organism level: while studying the onset or recovery of an infection, one might look at the probability that a single virion can infect a target cell and proliferate [9]; in epidemiology, the goal might be to estimate the probability that a newly introduced pathogen will become endemic in a na¨ıve host population [4]. The same multi-scale interest in invasions appears in the study of population genetics: at the multi-organism scale one studies the probability that a novel allele can fix in a population, possibly affecting the population’s overall fitness [17, 21, 25]; at the cellular level, the invasion of a mutation in a stem cell population has been studied as an important first step in certain cancers [27]. What distinguishes the dynamics at different scales are the state-dependent rates of events. While multi-organism models tend to feature movement and direct interaction among individuals [7], cellular dynamics may be more spatially static with indirect interactions (competition for resources, or signaling at a distance, for example) [29].

It is natural to model these population dynamics using continuous-time, discrete-state-space Markov chains. It is easy to encode nonlinear interactions through state-dependent transition rates and there are numerous straightforward simulation techniques available that can be exact, but costly [13], or inexact (with respect to boundary interactions), but efficient [3]. It is also straightforward to write down difference equations whose solutions represent the probability of invasion from a small number of individuals, or the mean extinction time starting from a large population size. Often it is even possible to find exact solutions for these systems of difference equations, but, importantly, it is difficult to interpret how these exact solutions depend on the parameters of the original system.

To overcome both computational and analytical challenges, it has become common to use Stochastic Differential Equations (SDEs) as a continuous-state-space approximation for the discrete-state-space Markov chains. Using Kurtz’s Diffusion Approximation Theorem as a guide [19, 2], there is a natural way to translate Markov chain transition rates into the drift and diffusion terms of an SDE. (See Section 3.1 for discussion.) This technique has been used prominently to model epidemics [1], neuronal activity [14, 10] and branching processes with logistic growth limits [20]. In the SDE setting, solutions to invasion probability and mean extinction time questions can be obtained by solving certain ODEs (or PDEs, depending on the dimension). These ODEs can also be obtained without appealing to an SDE model by applying a na¨ıve Taylor series approximation of the Kolmogorov Equations of the Markov chain model. (See Section 3.1 again for discussion.) However, multiple authors have shown that there are important differences between Diffusion Approximations and their corresponding CTMCs. For example, Doering and co-authors [11, 12] have shown that the mean extinction time for SDE models can differ substantially from that of the corresponding Markov chain model. Newby and Bressloff also documented the issue for neuronal models [16, 22, 8].

While this issue of the mean extinction time has received considerable recent attention, the probability of invasion question was studied, and in many ways resolved, much earlier. Frank Ball and colleagues [4, 5] showed that the initial phase of a certain class of epidemic models can be well-approximated by branching processes for which an exact probability of invasion can be calculated. For example, in the large population limit, the probability that a single infectious individual will cause an epidemic in the stochastic Susceptible-Infectious-Susceptible (SIS) and stochastic Susceptible-Infectious-Recovered (SIR) models is 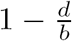, where *b* is the rate at which an infectious individual infects susceptible members of the population and *d* is the rate at which the pathogen is cleared from an infectious individual. (Throughout this work, we will use the letters *b* and *d* to evoke the words “birth” and “death” to correspond to population increases and decreases, regardless of the particular application.) This formula is appealing because it is trivial to understand the impact of model parameters, but we note that because this method relies on an infinite population size limit, it accordingly gives no guidance on how invasion probabilities scale with finite population sizes. Moreover, the Branching Process Approximation is only valid when the dynamics are “asymptotically linear” in a sense that will described after Assumption 1.1 below. Heuristically, this means that a “birth event” only depends on one member of the population of interest. This is often not the case in chemical reactions (see Doering, 2007 [12] for an example) and a recent investigation into the impact of sensing and decision-making on animal movement has revealed non-linear encounter rates in predator-prey models [15].

The above considerations motivate the present work, in which we analyze the performance of standard continuum approximations for the probability of invasion in Markov chain models. The mathematical framework allows for nonlinear transition rate functions and provides a natural way to estimate probability of invasion when the population size *N* is finite. We perform asymptotic analysis (*N →* ∞) on the Diffusion Approximation and a WKB-like Exponential Approximation, showing that the former yields an incorrect value in essentially all cases. When valid, the Exponential Approximation appears to yield the correct large *N* limit. Remarkably though, when the population size is small or intermediate, and when the parameters of the problem form a system that is near-critical, we show numerically that the Diffusion Approximation captures invasion probabilities much better than the Exponential Approximation. Rather than declare one approximation or another the “winner” we think it would be most useful for practitioners to keep in mind the tradeoffs involved in choosing which method to implement in their own work.

### 1.1. Mathematical Framework

For a given population scale *N*, consider a continuous-time Markov chain *X*_*N*_(*t*) that takes its values in the set of integers 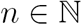. We consider the class of models with transitions of size one and the transition rates from the state *n* are given by

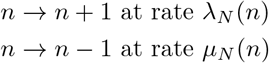

with 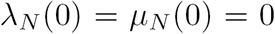. While 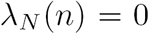 is allowable for some *n* > 0, we require that the death rate satisfies 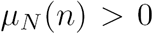 for all 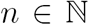. It follows that 0 is the only absorbing state.

ASSUMPTION 1.1 (Transition rate shape functions). *There exist functions* λ : 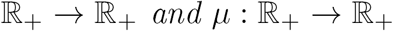 *such that*

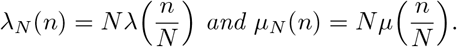

*Furthermore*,

a. *there exists an A* ∈ (0, ∞] *such that* λ *and µ are differentiable for all x* ∈ (0, *A*);
b. λ(0) = *µ*(0) = 0; *λ*(*x*) ≥ 0 *for all x* ∈ (0, ∞); µ(*x*) > 0 *for all x* ∈ (0, ∞); *and*
c. *there exists a value x_*_* > 0 *such that* λ(*x*_*_) = *µ*(*x*_*_) *and for all x* ∈ (0, *x*_*_), *either λ*(*x*) > *µ*(*x*) *or µ*(*x*) < *λ*(*x*). *In the former case, the system is called* supercritical; *in the latter*, subcritical.

We will refer to *x*_*_ as the *minimal rate-balanced point.* When the system is discrete, rate-balanced points are not always precisely achieved. For a given system size *N*, we define the discrete analogue *N*_*_ as follows.

DEFINITION 1.2. *For a given* 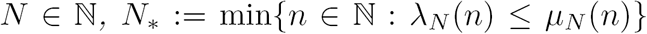 *for supercritical systems and* 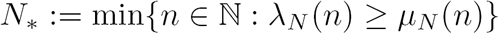 *for subcritical systems.*

If the transition rate shape functions are differentiable at zero with 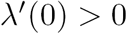 and 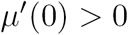, then we call the system *asymptotically linear.*

*Example* 1 (Stochastic epidemics). For a population size *N*, and constants *b* > 0 and *d* > 0, the size of the infectious population in a stochastic (non-density dependent) Susceptible-Infectious-Susceptible system is defined by the transition rates

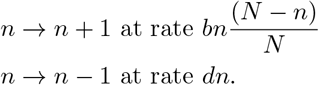

These dynamics are characterized by the rate shape functions

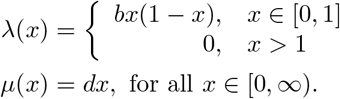

**Fig. 1.**
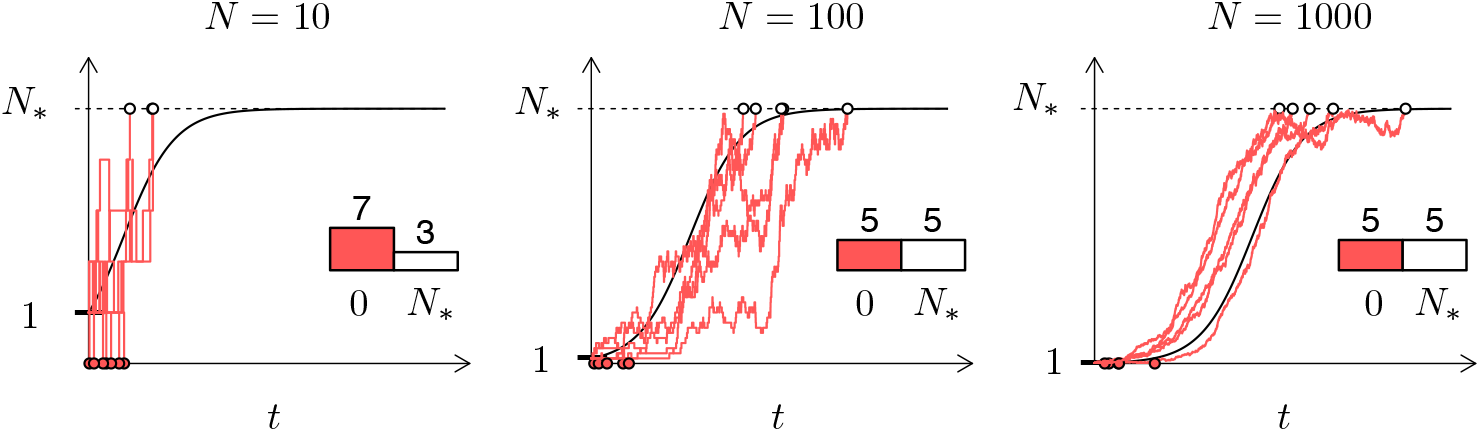
Number of infectious individuals resulting from the introduction of one infectious individual in a population of size N = 10, 100, and 1000. Ten sample paths for the stochastic SIS model defined by Example 1 are plotted (red lines) with the solution to the analogous ordinary differential equation model (black curve). Our representation for having achieved the endemic state is N_*_ (dashed horizontal black line), which is defined in Definition 1.2. Open circles are plotted when N_*_ is reached before 0 and red points indicate when the pathogen died out of the population before reaching N_*_. For all simulations b = 2 and d = 1.

The minimal rate-balanced point *x*_*_ = 1 – *d/b*, and for the process 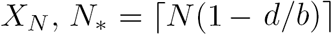.

*Example* 2 (Density-dependent mortality). In this case, the population scale *N* plays the role of the carrying capacity of an environment. Again, let *d* > 0 and let *b* > 0 be the growth rate of the population when scarcity of resources is not a factor. Then we consider the dynamics set by the rate shape functions

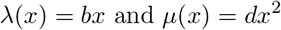

for *x* ∈ [0, ∞). We note that this is the standard logistic growth model with the intrinsic growth rate parameter *b = r* and death rate *d = r/K* for a carrying capacity *K.* The resulting transition rates are similar to the Logistic Branching Process presented by Amaury Lambert (2005) [20]. The rate-balanced value for this system is *x*_*_ = *b/d*, and for the process *X*_*N*_, we have 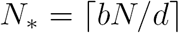

*Example* 3 (Resource-constrained birth). Let *b* > 0, *d* > 0 and *a* > 0. We define our dynamics according to the rate shape functions

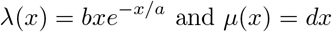

for *x ∈* [0, ∞). Note that 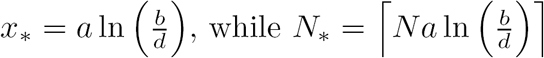.

*Example* 4 (General nonlinear single term model). Let *b, β, d, δ* > 0 with *β ≠ δ.* We define our dynamics according to the rate shape functions

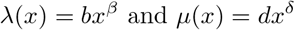

for *x ∈* [0, ∞). If *β < δ* we say that *λ* has the leading order. If *β > δ* we say that *µ* has the leading order. In this case, *x*_*_ = (*b*/*d*)*^β-δ^*, and for the process *X*_*N*_, 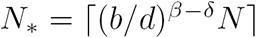.

While conducting our asymptotic analysis, we will often prefer an alternative assumption for the transition rate functions that features explicit series expansions for the birth and the death rates.

ASSUMPTION 1.3 (Series representation for the rate shape functions). *Let λ and µ be as in Assumption 1.1. Additionally, we assume that there exist constants b*, *β, d, δ*, > 0 *with* 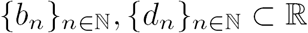, *and an integer* 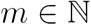, *such that*

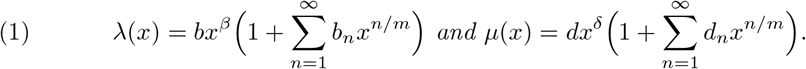

*for all* 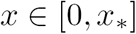.

For simplicity in the analysis, we also assume that *β* and *δ* are rational numbers and m is chosen such that *βm* and *δm* are integer values.

### 1.2. Approximations for Invasion Probabilities

The conditions of Assumption 1.1 ensure that as *N →* ∞, the stochastic processes *X*_*N*_(*t*) converge pathwise to an associated ODE. Kurtz [19] defines the sense of this convergence rigorously as follows: Let *λ* and *µ* be locally Lipschitz functions. Define 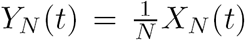, and suppose 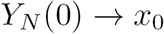. Define *x*(*t*) to be the solution to

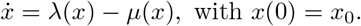

Then

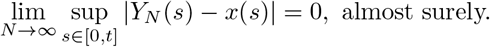

A visual demonstration of this result is shown in Figure 1 for the stochastic Susceptible-Infectious-Susceptible epidemic model (Example 1 above). Conditioned on not hitting zero, the solutions converge to the heteroclinic connection between zero and the minimal rate-balanced point *x*_*_. In our case, zero is an unstable fixed point of the limiting ODE. Thus, there is an apparent contradiction since, although a solution with initial condition zero should be zero for all *t ≥* 0, the fraction of solutions that avoid extinction *does not go to zero as N →* ∞. This conflict is resolved by noting that the theorem above only applies for a fixed time window. It is indeed true that for any fixed time 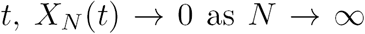. However, it is also true that, conditioned on successful invasion, the hitting time for *N*_*_ (Definition 1.2) tends to infinity as well.

We call the probability of hitting *N*_*_ before 0 the invasion probability and adopt the following notation.

DEFINITION 1.4. *For the process X_N_*(*t*), *we define the* hitting time *τ*_*N*_ *as follows:*

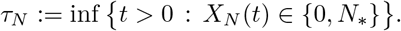

*We define the associated extinction probabilities* 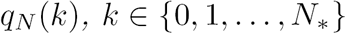 *by*

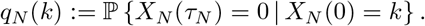

*From this we define the asymptotic probability of invasion p_invasion_*(*k*) *(conditioned on k individuals at time zero) to be*

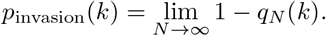

It is possible to obtain an exact solution for these invasion probabilities, and we provide a formula in Section 2. In 1983, Frank Ball [4] introduced a slightly different definition of an invasion probability (which he referred to as the *probability of a true epidemic*) and, in the asymptotically linear case, proved that the probability of a true epidemic converges to the survival probability of an associated branching process. To define this associated branching process, consider the number of offspring that any given individual gives birth to before expiring. In the asymptotically linear case, the leading order terms of the birth and death rates are *b* and *d*, respectively. Therefore, the number of births before the individual dies has a Geometric distribution with "success probability" *d*/(*b + d*).

DEFINITION 1.5 (Branching Process Approximation). *Suppose that the transition rate shape functions λ and μ, satisfy Assumptions 1.1 and 1.3 and that the leading order exponents satisfy β = δ* = 1. *Define the discrete time stochastic process* 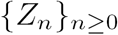 *to be Z*_0_ = *k and for* 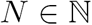,

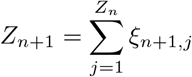

*where* 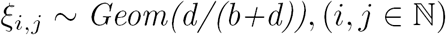 *are independent and identically distributed. Then, we define*

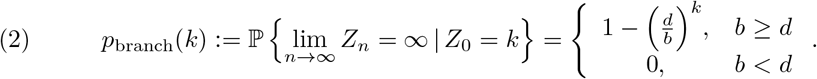

Ball [4] proved that *p*_invasion_(*k*) = *P*_branch_(*k*) in this setting. What the Branching Process Approximation lacks is a clear interpretation in asymptotically nonlinear cases (e.g. Example 4) and a means to consider what the invasion probability is for finite *N*. For these cases we consider the Diffusion Approximation and the Exponential Approximation. The derivations of these approximations are given in Sections 3 and 4, respectively. In what follows we introduce the small parameter ε, which corresponds to the inverse of the population scale parameter *N*.

**Table 1.**
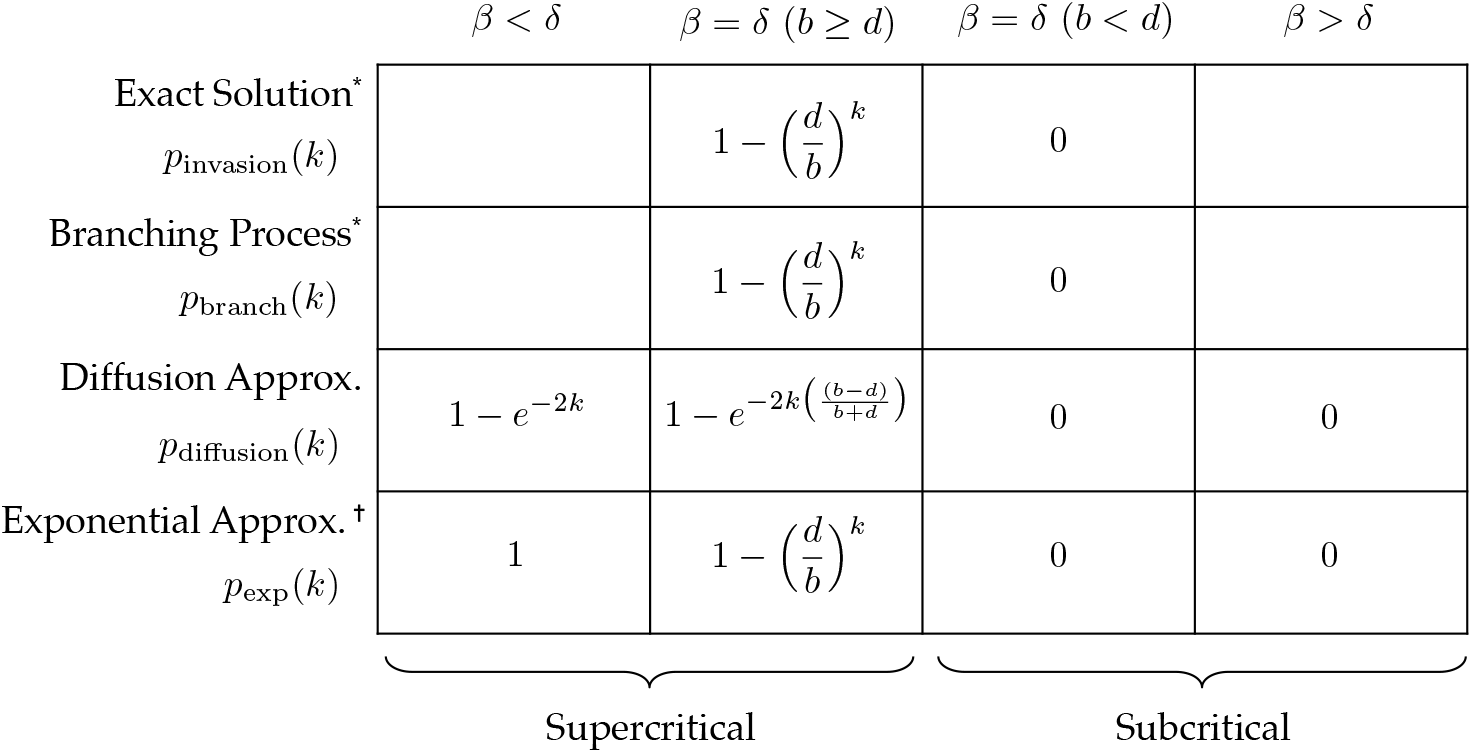
Summary of asymptotic invasion probabilities (see Definition 1.4). * These results are only proved for the asymptotically linear case (i.e. when β = δ = 1). ^†^We define the Exponential Approximation to be zero for subcritical regimes because the asymptotic limit tends to zero, but the approach to zero is through negative values.

| | $\beta < \delta$ | $\beta = \delta \ (b \geq d)$ | $\beta = \delta \ (b < d)$ | $\beta > \delta$ |
| --- | --- | --- | --- | --- |
| Exact Solution*<br>$p_{\text{invasion}}(k)$ | | $1 - \left(\frac{d}{b}\right)^k$ | 0 | |
| Branching Process*<br>$p_{\text{branch}}(k)$ | | $1 - \left(\frac{d}{b}\right)^k$ | 0 | |
| Diffusion Approx.<br>$p_{\text{diffusion}}(k)$ | $1 - e^{-2k}$ | $1 - e^{-2k\left(\frac{b-d}{b+d}\right)}$ | 0 | 0 |
| Exponential Approx. <sup>†</sup><br>$p_{\text{exp}}(k)$ | 1 | $1 - \left(\frac{d}{b}\right)^k$ | 0 | 0 |

Supercritical Subcritical

DEFINITION 1.6 (Diffusion Approximation). *Let ɛ* > 0 *be given. Then for x ∈* 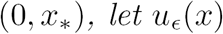 *be the solution to the boundary value problem*

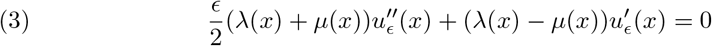

*with* 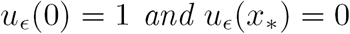.

*Then, setting* ∈ = 1/*N, we define the* Diffusion Approximation *for invasion probabilities to be the relationship*

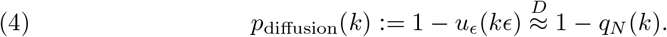

DEFINITION 1.7 (Exponential Approximation). *Let ɛ* > 0 *be given. Then for* 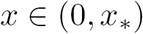, *define* 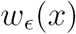 *to be*

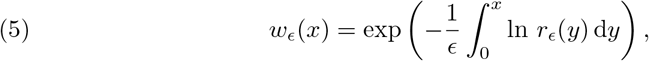

*where r*_*ɛ*_(*x*) *satisfies*

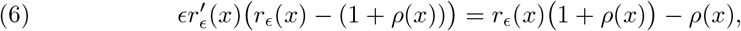

*with ρ*(*x) = λ*(*x*)/*µ*(*x*). *Then the* Exponential Approximation *for invasion probabilities is defined by the relationship*

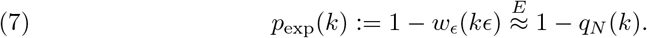

In Table 1 we summarize the results in Sections 3 and 4. Consistent with asymptotic analyses of the Diffusion Approximation for corresponding extinction probabilities, we find that the Diffusion Approximation disagrees with the Branching Process Approximation in the large *N* limit. Remarkably when the leading orders of the transition rate shape functions *λ* and *µ* do not match, the Diffusion Approximation reduces to two cases, ignoring all detailed information contained in the rate functions. In contrast, the Exponential Approximation does provide the right large *N* limit. We note however, that the Exponential Approximation is only well-defined in the supercritical case.

In Section 5 we investigate the problem numerically and find mixed results. In the large *N* limit, the Exponential Approximation agrees with the Markov chain model, even in the asymptotically nonlinear cases we consider. However, when the population size parameter *N* is of small or intermediate size, say *N* = 50, it is common that the Diffusion Approximation more faithfully represents the Markov chain model, especially when the parameter set is near-critical.

## 2. Exact solution for invasion probabilities

PROPOSITION 2.1. *For fixed* 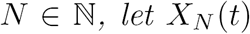 *be a CTMC with transition rate shape functions λ and µ as defined in Assumption 1.1. Let* 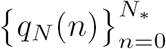 *be the extinction probabilities as defined in Definition 1.4. Then the collection* 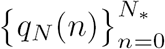 *satisfies the system of difference equations*

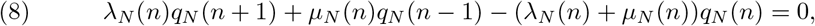

*with* 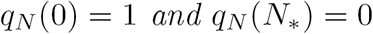

*Proof.* Though the proof appears in standard texts, we include it for completeness. Fix 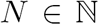 and let an initial condition 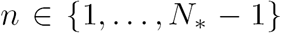 be given. Define *T :=* inf{*t* > 0 : *X*_*N*_(*t*) ≠ *n*}, the first time the process changes states. Then, applying the Strong Markov property, we have

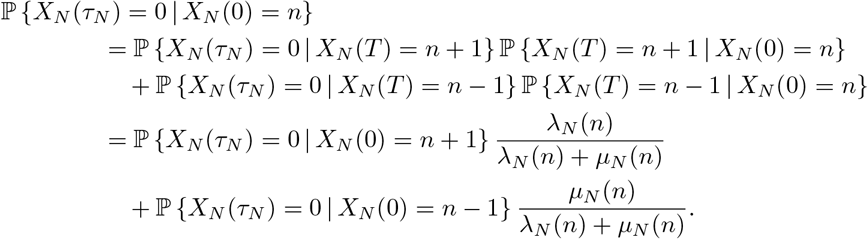

Applying the *q*_*N*_(*n*) notation from Definition 1.4 and multiplying through by *λ*_*N*_(*n*) + *µ*_*N*_(*n*), we have (8), where 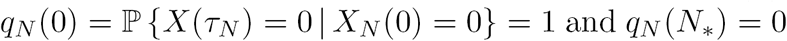.

The following argument for finding the exact solution is similar to one presented on birth-death processes in Norris [23], for example, and has a similar form to the exact solutions for mean extinction times presented by Doering et al [11, 12].

Proposition 2.2 (Exact solution for extinction probability). *Let X*_*N*_(*t*) *be defined as in Proposition 2.1, with associated extinction probabilities* 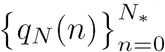. *Then* for *k* ∈{1, 2, …, *N*_*_−1},

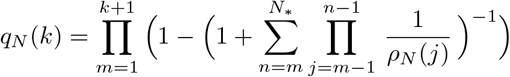

*where we define* 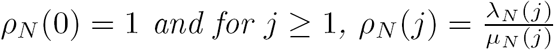.

*Proof.* Suppressing the dependence on *N*, we write

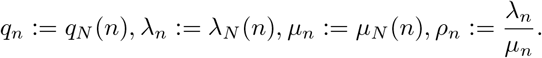

Rearranging (8), we have

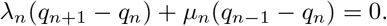

Introducing *δ*_*n*_ := *q*_*n*−1_ − *q*_*n*_, we can solve for *δ*_*n*+1_ and extrapolate to a dependence on *δ*_*k*_:

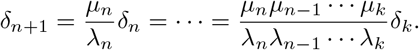

Recalling Definition 1.4, we have the boundary conditions 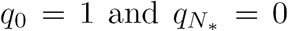. It follows that

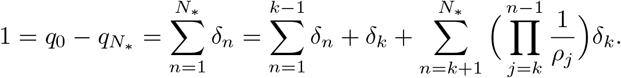

Noticing that 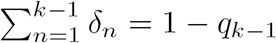, we have

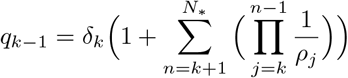

and solving for *δ*_*k*_

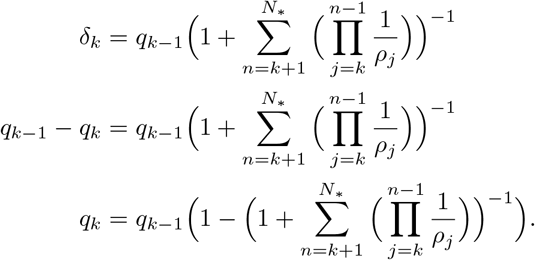

From this recursion we obtain

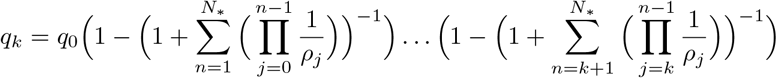

and since *q*_0_ = 1, the result follows.

## 3. Diffusion Approximation

## 3.1. Motivations for the Diffusion Approximation

There are two points of view on the derivation of the Diffusion Approximation for invasion probabilities. They yield the same result. The first point of view (Motivation 1, below) is common in the physics literature [6]. It involves substituting a smooth function *u*_*∈*_ into the difference equation (8) and then converting it to a differential equation by writing out Taylor expansions and matching terms. In a second point of view (Motivation 2, below), we define a Stochastic Differential Equation whose infinitesimal first and second moments are defined to match that of the original birth-death chain in a sense discussed in [26] and [2]. Then the same Diffusion Approximation results from computing the probability that this SDE hits the rate-balanced state before going extinct, starting from the value *k*_*ɛ*_ = *k*/*N*, where *k* is the number of initially introduced individuals.

### Motivation 1: Taylor series approximation

To begin this approach, we assume that *u*(*x*) is a smooth function and appeal to the following formal argument. We start with the difference equation

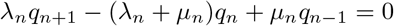

and rewrite it using the rate shape functions according to Assumption 1.1, *λ*_*n*_ = *Nλ*(*n/N*) and *µ*_*n*_ = *Nµ*(*n/N*), along with *q*_*n*_ = *u*(*n/N*):

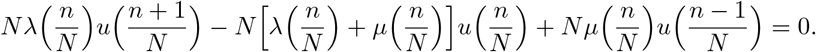

Writing *x = n/N* and ∈ = 1/*N*,

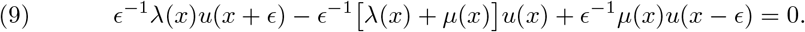

We then apply a Taylor series approximation to the *u*(*x* + *ϵ*) and *u*(*x* - *ϵ*) terms:

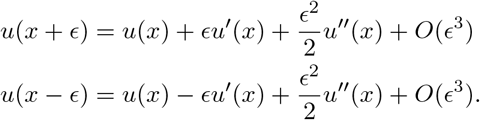

Substituting these into (9) and neglecting the higher order terms, we have

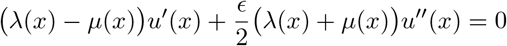

which is the form seen in Definition 1.6.

*Motivation 2: Hitting probabilities for an SDE approximation*

Let *ϵ* = 1/*N* and *y = n/N*, and consider the following calculation for the infinitesimal mean:

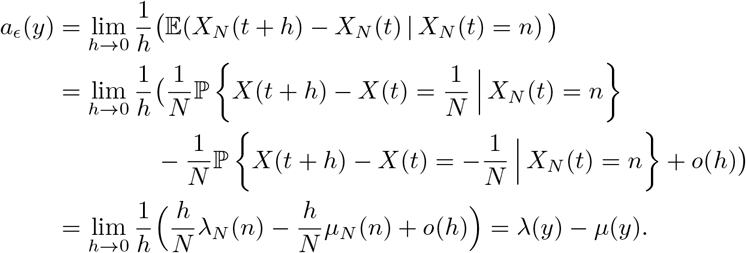

In the last equality we used Assumption 1.1: *λ*_*N*_(*n*) = *Nλ*(*n/N*) and *µ*_*N*_(*n*) = *N µ*(*n/N*). Note that the infinitesimal mean does not, in fact, depend on *N*. On the other hand, a factor of 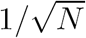 does appear in the calculation of the infinitesimal second moment:

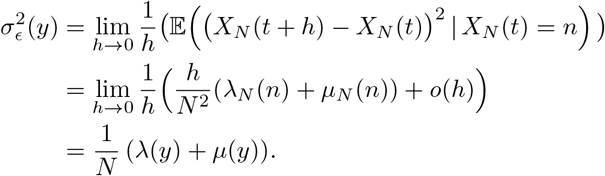

Using these values, we define the SDE approximation as follows.

DEFINITION 3.1 (SDE Approximation). *Let ϵ* > 0. *We say that a process Y_ϵ_*(*t*) *is an* SDE approximation *for the family of CTMCs defined with rate shape functions* λ *and µ if it solves the Itˆo Stochastic Differential Equation*:

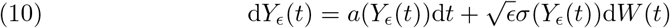

where

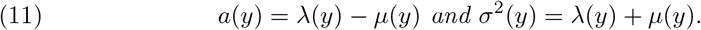

*Specifically, if ϵ* = 1/*N, then the SDE associated with the CTMC X_N_*(*t*) *is Y*_*ϵ*_(*t*).

PROPOSITION 3.2. *Let Y*_*ϵ*_(*t*) *be defined as in* (10) *and* (11). *Let T*_*ϵ*_ := inf{t > 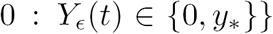. *Then* 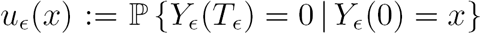 *satisfies the boundary value problem* (3).

*Proof.* The proof follows quickly from stochastic calculus. For a general continuous sample-path SDE with infinitesimal generator *L*, the probability *u*(*y*) that the process hits a value *a* before *b* starting from *y* ∈ [*a*, *b*] satisfies the boundary value problem

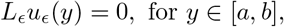

with *u*(*a*) = 1 and *u*(*b*)=0 [18].

It remains to note that the generator of *Y*_*ϵ*_(*t*) is

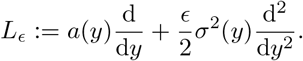

## 3.2. Analysis of p_diffusion_(k). It is useful to introduce the functions

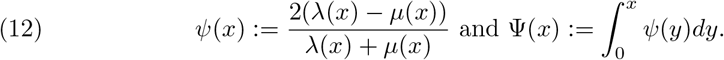

We note that *ψ*(*x*) in (12) can be expressed exclusively in terms of the ratio *ρ*(*x*) := *λ*(*x*)/*µ*(*x*) as

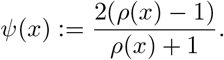

However, in order to retain the ability to directly interpret results in terms of explicit birth and death rate parameters, we generally perform our analyses and present our results using the rate shape functions λ and *µ* directly, as they are characterized in Assumption 1.1. An exception arises in Section 4, where we find that it is most natural to analyze the Exponential Approximation in terms of the ratio of rate functions *ρ*(*x*).

PROPOSITION 3.3. *Let u*_*ϵ*_*(x) be the Diffusion Approximation for the extinction probabilities, satisfying* (3) *of Definition 1.6. Then*

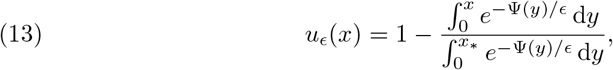

*where x*_*_ = min{*x* : *λ*(*x) = µ*(*x*)}.

*Proof.* It we let *h*(*x*) = 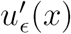 then the ODE (3) becomes

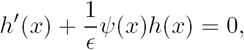

which has general solution 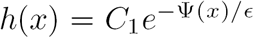. Integrating to get 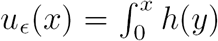 d*y* and enforcing the boundary conditions *u*_ϵ_(0) = 1 and *u*_*ϵ*_(*x*_*_) = 1 yields the solution (13).

We note that a similar formula can be found in Pakmadan et al. [24].

We summarize the asymptotic behavior of the Diffusion Approximation as follows.

THEOREM 3.4. *Suppose that the transition rate shape functions* λ *and µ each admit a series expansion as defined by Assumption 1.3. If the leading order exponents β and δ are equal, then for 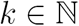*,

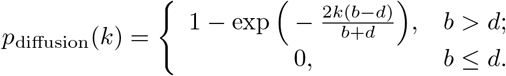

*Otherwise, if β ≠ δ, then*

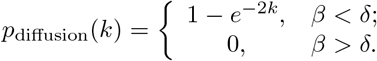

Remarkably, the Diffusion Approximation essentially ignores the detailed structure of the rate functions when they have different leading orders.

The integrals that appear in (13) are in a form compatible with applying Watson’s Lemma [28]. In fact, Watson’s Lemma can be applied directly to the denominator. The subtlety in the present analysis is that, in the numerator, the upper limit of integration depends on *ϵ.* To proceed, we follow Laplace’s method and introduce a substitution. Suppose first that we are in the supercritical case, as defined by Assumption 1.1, so that for all *x* ∈ (0, *x*\*) we have that *ψ*(*x*) > 0. It follows that Ψ(*x*) is increasing on that interval and therefore invertible. Letting *t* = Ψ(*y*) we have

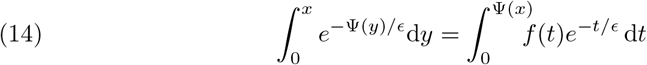

where

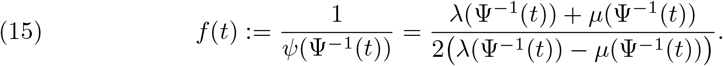

On the other hand, in the subcritical case, *ψ*(*x*) < 0 when *x ∈* (0, *x*\*). Introducing the notation *ψ_r_*(*x*) = − *ψ*(*x*) and Ψ_*r*_(*x*) = − Ψ(*x*), we substitute *t* = Ψ_*r*_(*y*) to attain

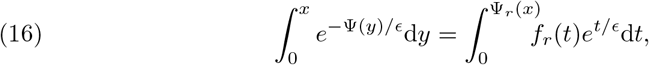

noting that the exponential in the second integral has a positive exponent, where

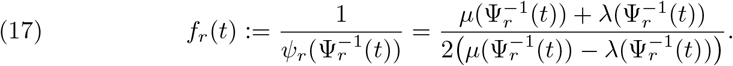

The proof follows from a combination of three lemmas. In Lemma 3.7 we assume that *f*(*t*) admits a series expansion and then perform an asymptotic analysis on the ratios that appear in integrals of the form (14) and (16). In Lemmas 3.5 and 3.6 we establish that *ψ*(*x*) and *f*(*t*) admit series expansions when *λ*(*x*) and *µ*(*x*) do, and provide the coefficients and powers of the first few terms. After presenting these lemmas and their proofs, we complete the proof of Theorem 3.4.

LEMMA 3.5. *Suppose that the transition rate shape functions λ and µ satisfy Assumption 1.3. Then there exists an expansion for ψ*(*x) of the form*

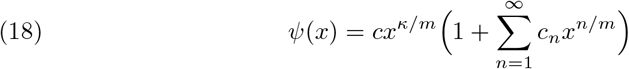

*where*

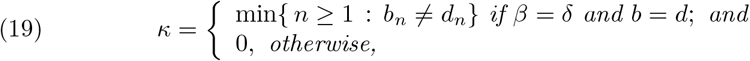

*and the value of c is given in the following table.*

| $c$ | $\beta < \delta$ | $\beta = \delta$ | $\beta > \delta$ |
| --- | --- | --- | --- |
| $b \neq d$ | 2 | $\frac{2(b-d)}{b+d}$ | -2 |
| $b = d$ | 2 | $(b_\kappa - d_\kappa)$ | -2 |

*Proof.* Substituting the expansions (1) in for *λ* and *µ*, we have

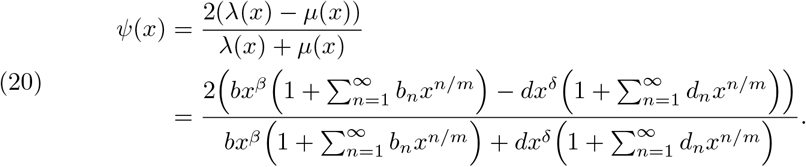

*Case 1: β = δ, b ≠ d.* We can factor out *x*^*β*^ = *x*^*δ*^ from the numerator and denominator. Then (20) simplifies to

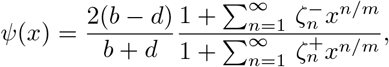

where 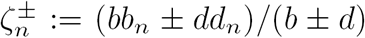. Following the notation introduced in Proposition A.2 of the Appendix, we have

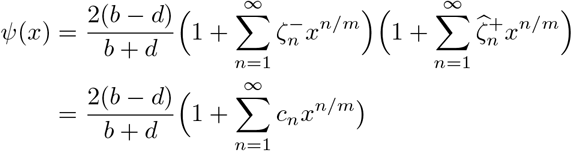

so that in the notation of (18), we have *c* = 2(*b-d*)/(*b + d*) and *κ* = 0. The remaining coefficients can be expressed in terms of 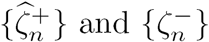:

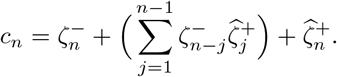

*Case 2: β = δ, b = d.*

In this case we can factor *bx*^*β*^ = *dx*^*δ*^ from the numerator and denominator of (20), which leaves

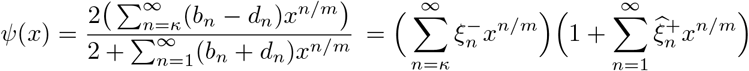

where 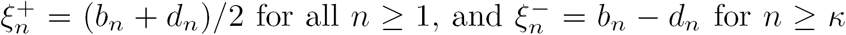. The form of 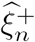 follows from Proposition A.2 in the Appendix. It follows that *ψ*(*x*) can be written in the form of (18) with the leading order exponent being *κ*/*m* and

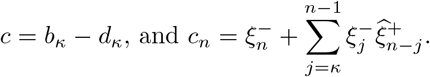

*Case 3: β < δ*

In this case we begin by factoring out *bx*^*β*^ from the numerator and denominator of equation (20). After cancellation we have

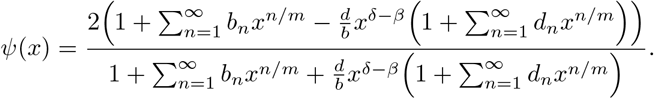

From this form and Proposition A.2, we see that *κ* = 0 and *c* = 2. Furthermore, the existence of the sequence *c*_*n*_ follows from our assumption that (*β* - *δ*)*m* is an integer.

*Case 4: β > δ*

In this case we begin by factoring out *dx*^*δ*^ from the numerator and denominator of equation (20). After cancellation and some rearrangement we have

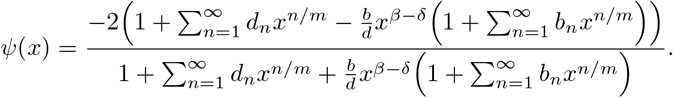

Similar to Case 3, we see that *κ* = 0 and *c* = −2.

From here, the proof proceeds in two steps. First we need to rewrite the integral Ψ(*x*) presented in (12) in a form that is compatible with Lemma 3.7.

LEMMA 3.6. *Suppose that the shape functions λ and µ satisfy Assumption 1.3 and we write ψ as described in Lemma 3.5. Furthermore, let* 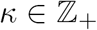 *be as defined in Lemma 3.5. Then f(t) admits a series expansion of the form*

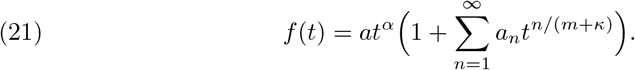

*where m is the positive integer defined in Assumption 1.3 and*

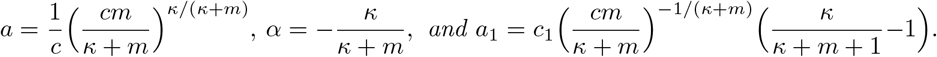

*Proof.* It will be convenient to write the series expansion for *ψ* in the form

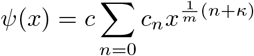

with the convention that *c*_0_ = 1. For the main part of the proof, we will take the *c* > 0 case. At the end, we will explain how the argument changes when *c* < 0. Integrating term-by-term, we have

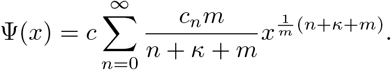

It will be useful to define *h*(*x*) := Ψ(*x*^*m*^) with a series expansion written as

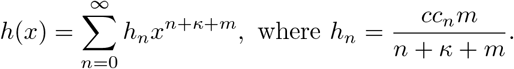

Since *c* > 0, Ψ (*x*) is increasing on [0, *x*_*_) and therefore *h*(*x*) is increasing on 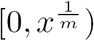. Setting *t = h*(*x*), by Proposition A.3, we can express the inverse of *h* in terms of the series

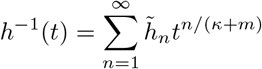

where

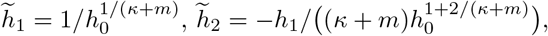

and so on.

We now turn our attention to the integral of interest, introducing the substitution 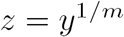, we have

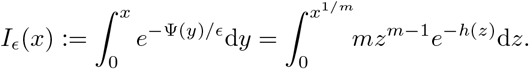

Then, taking *t* = *h*(*z*), and observing that 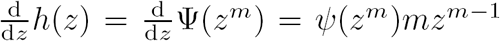, we have

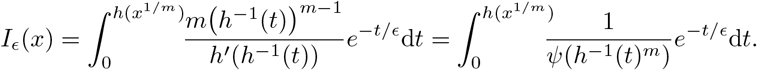

To determine the series expansion for *f*(*t*) := 1 /*ψ* (*h*^−1^ (*t*)^m^), we need to analyze the expansion for the denominator:

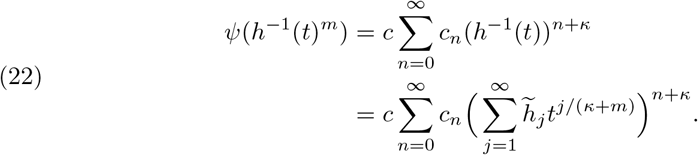

It is helpful to rewrite *h*^−1^(*t*) as follows

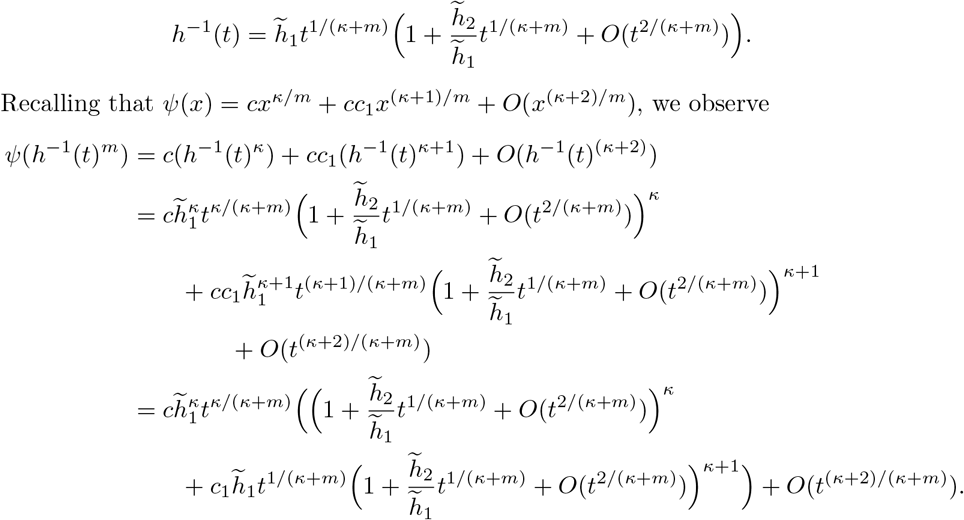

Then by applying Proposition A.1 twice, we obtain

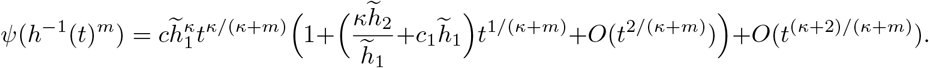

Proposition A.2 then yields

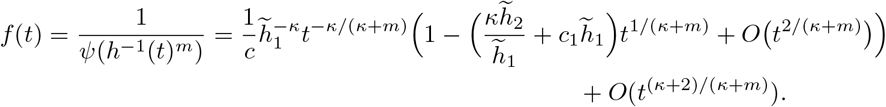

After simplifying, the stated results for *a, α* and *a*_1_ hold.

When *c* < 0, we replace *ψ* and Ψ with *ψ*_*r*_ and Ψ_*r*_ respectively. The leading term of *ψ*_*r*_ is then *-c*, which is positive and exact same procedure holds.

LEMMA 3.7. *Suppose that* 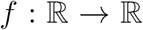 *admits a series expansion of the form* (21) *with* 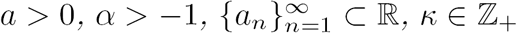 *and* 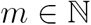. *Furthermore, let g* : 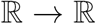 *be a continuously differentiable function that is monotonically increasing or decreasing for all* 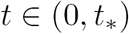, *with g*(0) = 0. *Define*

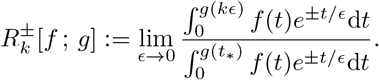

*Then*

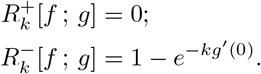

*Proof.* Our method is to replace *f*(*t*) with its series expansion and consider the integration term-by-term. Our results will be expressed in terms of the incomplete gamma function

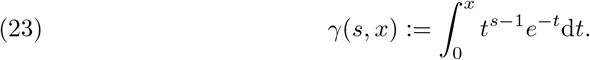

Near zero, the incomplete gamma function has the behavior that

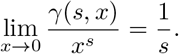

As usual, we define 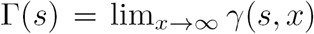. Note that for any *s* > −1 and *A* > 0, under the substitution *y* = *t/ϵ*, we have

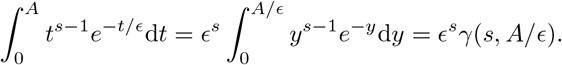

Therefore,

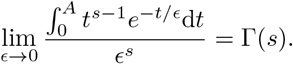

Writing *G* for *g*(*kϵ*) and *g*(*t*_*_) respectively, it follows that the numerator and the denominator appearing in 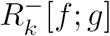 have the form

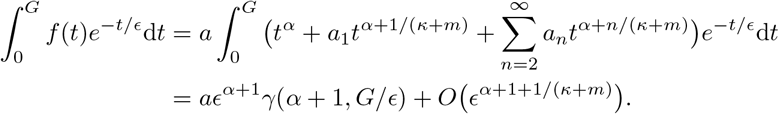

In taking the ratio, the coefficients 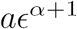 cancel, and neglecting higher order terms, we have

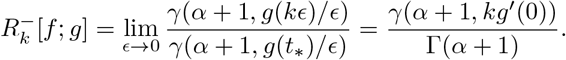

When *α* = 0, this takes the form

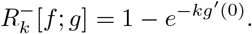

For the plus-sign case, we have

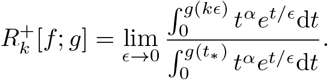

Since 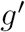 is continuous, let 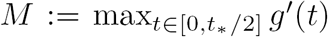. Then by the Taylor Remainder Theorem, since *g*(0) = 0 we have

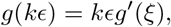

for some 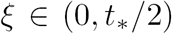. It follows that 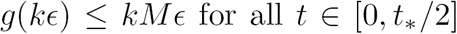. So for the numerator we have

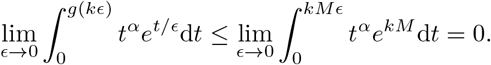

To study the denominator, we again make the substitution *y* = *t*/*ϵ* and obtain

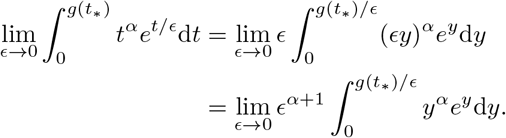

Noticing that the integral on the right hand side diverges to infinity, we rewrite the expression and apply l’Hôpital’s rule to find that the denominator diverges to infinity as ϵ → 0. It follows that the ratio tends to 0 as *ϵ →* 0.

We introduced the notation *f* and *g* to emphasize that the contribution to the final value comes from the limit of integration, not the function *f* in the integrand.

*Proof of Theorem 3.4.* First, recalling Definition 1.6 and the result from Proposition 13, we find that

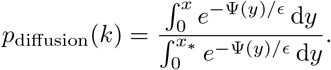

Noticing the familiar form of this expression, we apply Lemma 3.6 with *g* set to Ψ to rewrite p_diffusion_(*k*) in a form compatible with Lemma 3.7.

When *β >δ* or *β = δ* with *b ≤ d* Lemma 3.7 implies

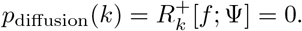

Alternatively, if *β* < *δ* or *β* = *δ* with *b > d*, Lemma 3.7 shows that

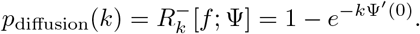

Since 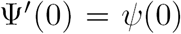, by Definition 12, the final result is attained by evaluating *ψ*(0) using Lemma 3.5.

## 4. Exponential approximation

Doering et al [11, 12] demonstrated that making a WKB-type ansatz of the form 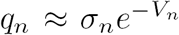 for some functions *σ* and *V*, can be an accurate method for constructing a continuum approximation for solving Kolmogorov equations. In the motivation that follows we provide a different analytical justification for this observation than has been presented elsewhere. We do this by transforming the system of equations defined by (8) into an equation for ratios instead of differences, then applying a Taylor series expansion technique similar to what is presented as Motivation 2 for the Diffusion Approximation.

## 4.1. Motivation for the Exponential Approximation

For *n* = 1, 2,…, *N*_*_, define

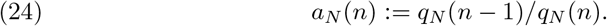

Since *q*_*N*_(0) = 1, we can write

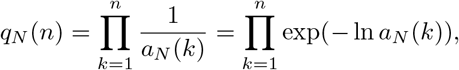

where

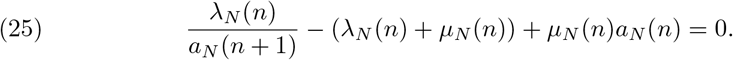

We can rewrite the exponent on the right-hand side to look like a Riemann sum:

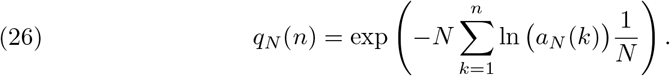

We note that the above form of the extinction probability is identical to that of the mean extinction time found in Doering et al. (2007) [12], where in our case 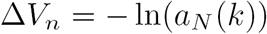. Mimicking the continuum approximations and Taylor expansions presented as motivation for the Diffusion Approximation, we introduce the function *r*_*ϵ*_(*x*) defined below. For *x = n*/*N* and *ϵ* = 1/*N*, we will write

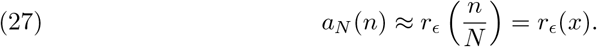

Then,

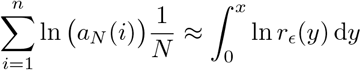

and the heuristic assertion is that (26) becomes

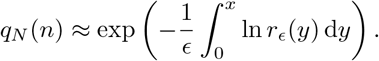

We note that this is the same form as presented in Definition 1.7 but it remains to characterize the function *r*_*ϵ*_(*x*). Indeed, following a Taylor series expansion technique, we will show that *r_ϵ_*(*x*) should have the form

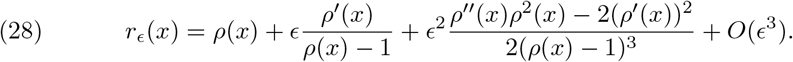

Dividing (25) through by *µ*_*N*_(*n*) and multiplying by *a*_*N*_(*n +* 1) we have

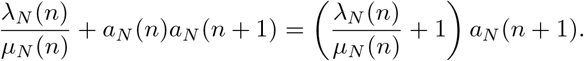

Let

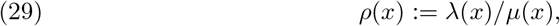

then using the definition of the transition rate shape functions *λ* and *µ*, we have

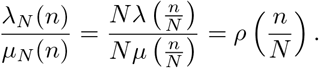

Thus,

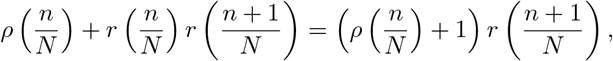

and, substituting *x* and *ϵ* for *n/N* and 1/*N*, we arrive at

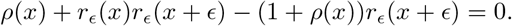

Neglecting higher order terms, we make the substitution

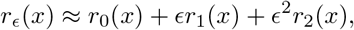

and obtain

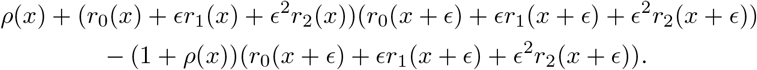

Now we perform a Taylor expansion on the *r*_*0*_(*x + ϵ*), *r*_1_(*x + ϵ*), and *r*_2_(*x* + *ϵ*) terms. Organizing the terms by powers of *ϵ*, we find the following system of equations:

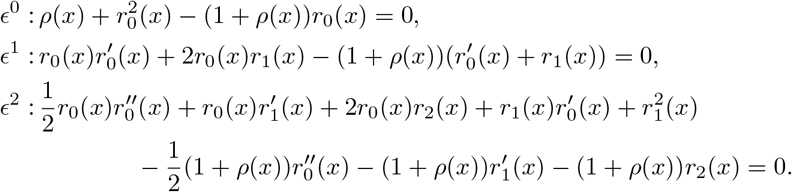

The first of these equations yields two solutions for *r*_0_(*x*): *r*_0_(*x*) = *ρ*(*x*) and *r*_0_(*x*) = 1. To show that the former must be true, consider that if *r*_0_(*x*) ≡ 1, then 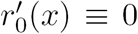. Substituting into the *ϵ*^1^ equation, it follows that *r*_1_(*x*) = 0. Continuing in this way, we find *r*_*i*_(*x*) = 0 for all *i* ≥ 1. This corresponds to the solution *q*_*N*_(*n*) = 1 for all *n* ∈ {0,…,*N*_*_}, which does not satisfy the boundary condition *q*_*N*_(*N_*_*) = 0. Therefore, we adopt the solution *r*_0_(*x*) = *ρ*(*x*). Feeding this into the order *ϵ*^2^ equation allows us to solve for *r*_1_(*x*). Similarly, substituting these solutions into the *ϵ*^2^ equation yields a solution for *r*_2_(*x*) and so on:

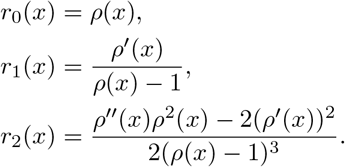

## 4.2. Analysis of *p*_exp_(*k*)

For the analysis that follows, we define what we call an *nth order* exponential approximation. Let

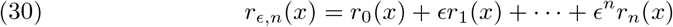

where the functions *r*_*j*_(*x*) are defined according to the procedure given in the preceding section. Then, we define

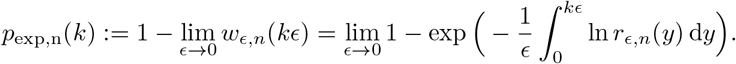

THEOREM 4.1 (Leading Order Exponential Approximation for Supercritical Systems). *Suppose that the transition rate shape functions λ and µ form a supercritical system, as defined in Assumption 1.1. Further suppose that* λ *and µ each admit a series expansion, as defined by Assumption 1.3.*

*If the leading order exponents β and δ are equal, then for any* 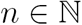,

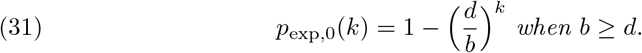

*Otherwise, when β < δ, p*_exp,0_(*k*) = 1.

*Remark* 4.2. When λ and *µ* form a subcritical system this method of approximation is not well defined. In this regime, the function 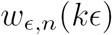 is often greater than one, meaning that 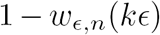 is negative and not a reasonable estimate for a probability. For example, note that this is what happens to equation (31) when *β = δ* and *b < d.* The proposed value for *p*_invasion_(*k*) is negative and cannot be an invasion probability. A practitioner could take the Exponential Approximation to be zero whenever the system of study is subcritical.

The proof of Theorem 4.1 is an immediate results of the following lemma.

LEMMA 4.3. *Suppose that the shape functions* λ *and µ each admit a series expansion as defined by Assumption 1.3, then 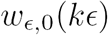 has the form:*

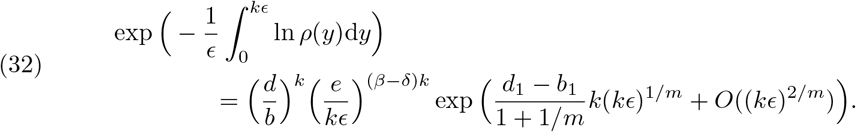

*Proof* First we find a series expansion for *ρ*(*x*). From the definition of *ρ* in equation (29) and the series expansions for *λ* and *µ* assumed in Assumption 1.3, we have

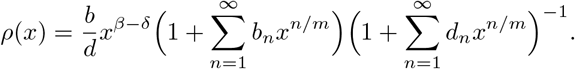

By applying Proposition A.2 and separating out the leading order terms, we obtain

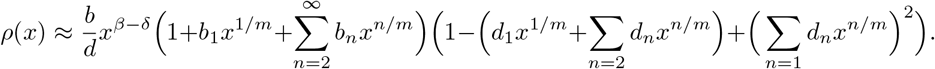

Collecting ordered terms yields the expansion

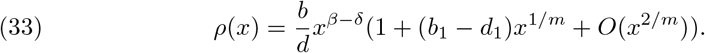

To consider the leading order approximation we use one term of 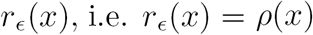. Then by plugging equation (33) into Definition 1.7, we have:

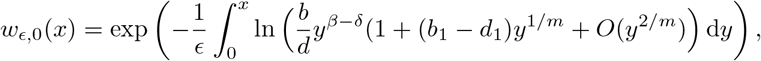

which can then be expressed in terms of having *k* individuals introduced

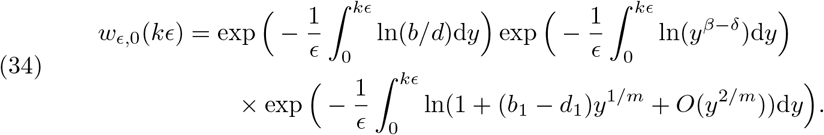

Considering the first term in the product of equation (34),

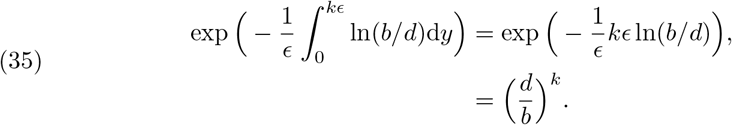

Then using an expansion for ln(1 + *x*) and integrating the result, the third term in the product becomes

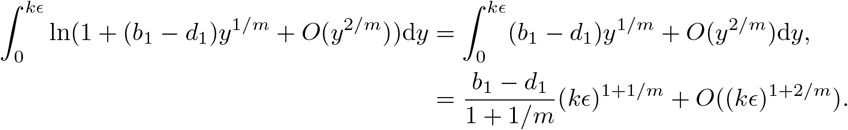

Thus the third term in the product of equation (34) can be rewritten as

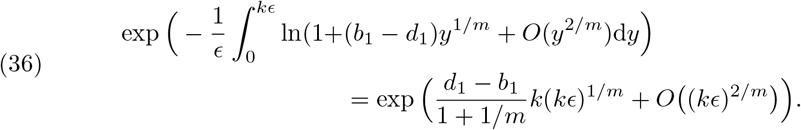

Rearranging and integrating the exponent from the second term in equation (34), we have

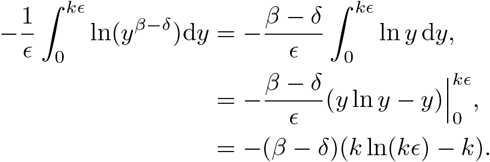

Therefore, the second term becomes

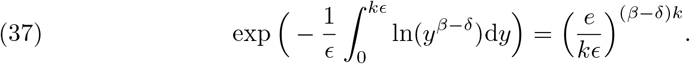

The result follows by rewriting equation (34) using equations (35), (36), and (37).

In Lemma 4.3, we show that the the Exponential Approximation exhibits a non-trivial dependence on *k*, the number of individuals introduced in the population. This dependence results from the relationship between the number of introduced individuals and the population scale *N*. We explicitly included the number of individuals introduced as a fraction of the population (*kϵ*) in the error term to record that error terms do not necessarily approach zero with *ϵ* if the number of individuals introduced is also allowed to vary.

To conclude our analysis, we return our attention to the functions *r*_*n*_(*x*) contained within equation (30). While we do not currently have a complete result for analyzing higher-order exponential approximations *p*_exp,*n*_(*k*) with *n* > 0, we would like to report the following recursion formula that can be used to generate these terms.

PROPOSITION 4.4. *For n ≥* 2, *r*_*n*_(*x*) *has the form*

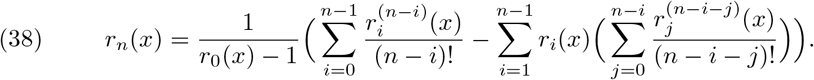

Proof. We begin by recalling the equation

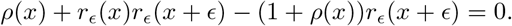

We then rewrite *r*_*ϵ*_(*x*) as a series in powers of *ϵ*

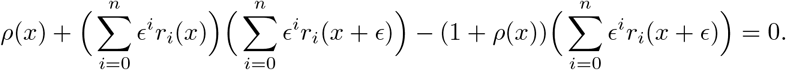

Substituting in Taylor expansions for the *r*_*i*_(*x + ϵ*) terms we obtain

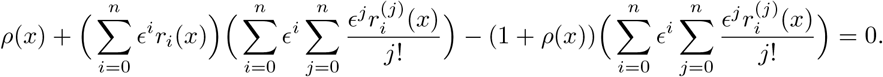

Since *r*_0_(*x*) = *ρ*(*x*) this reduces to

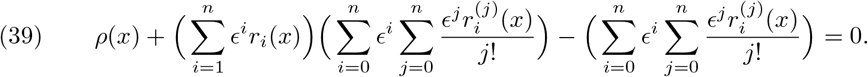

To solve for *r*_*n*_(*x*) we will consider the *ϵ*^*n*^ terms. First we consider the term

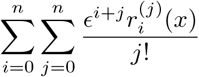

and notice that the *ϵ*^*n*^ terms are contained in the series

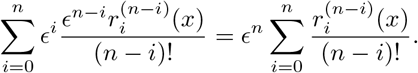

Now considering the second term in (39), we proceed by isolating the ϵ^*n-i*^ terms from the double summation in that product

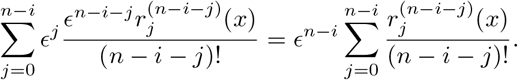

Thus, the *ϵ*^*n*^ terms in (39) can be expressed as

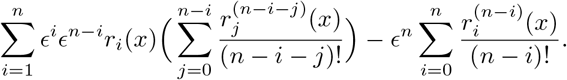

It follows from (39) that

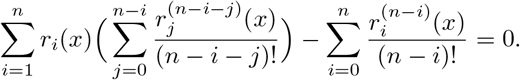

Then for the first term we have

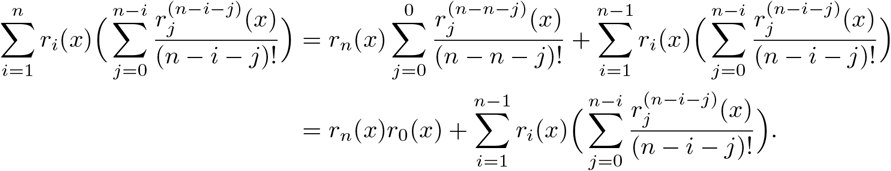

After also separating out the *r*_*n*_(*x*) term from the second series we have

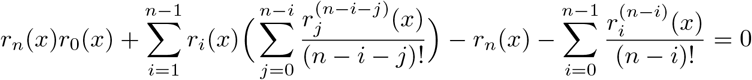

and

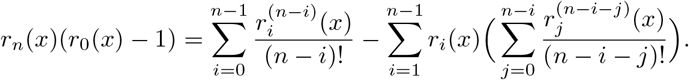

The final result is obtained by solving for *r*_*n*_(*x*).

## 5. Numerical observations for invasion probability approximations

In the preceding sections, our main analytical results for the Branching Process, Diffusion, and Exponential Approximations focus on the large population (*N* → ∞) limit. In this section, we validate our asymptotic results numerically and then shift our attention to consider the behavior of the approximations when the population size is small or intermediate. We challenge the methods in their ability to best approximate the exact solution as defined and calculated in Section 2. We focus our investigation on the specific examples presented in the introduction, and obtain exact solutions for the approximations whenever possible. We use numerical integration when it is not possible to obtain an exact solution for an approximation. In this way, we verify our analytical results and explore how the conditions of a system (e.g. population size and subcritical vs. supercritical dynamics) have an impact on whether a particular approximation method should be deemed fit for use. We set the death rate parameter (*d*) to be one by default.

### 5.1. Diffusion approximation methods fail for large populations when dynamics are supercritical

To complement our analysis in the previous section, we used exact solutions when possible and numerical integration when necessary (Simpson’s method coded in R) to evaluate the Diffusion Approximation and Exponential Approximation for invasion probabilities. In Figure 2, the results are shown for Examples 1, 2, and 3 for a range of population sizes and parameter value choices. We chose the exponents and coefficients so that the dynamics are supercritical and the Exponential Approximation is well defined. In each panel, we highlight that as the total population size becomes large, the values calculated numerically approach their corresponding analytically determined limit (indicated by “*”).

**Fig. 2.**
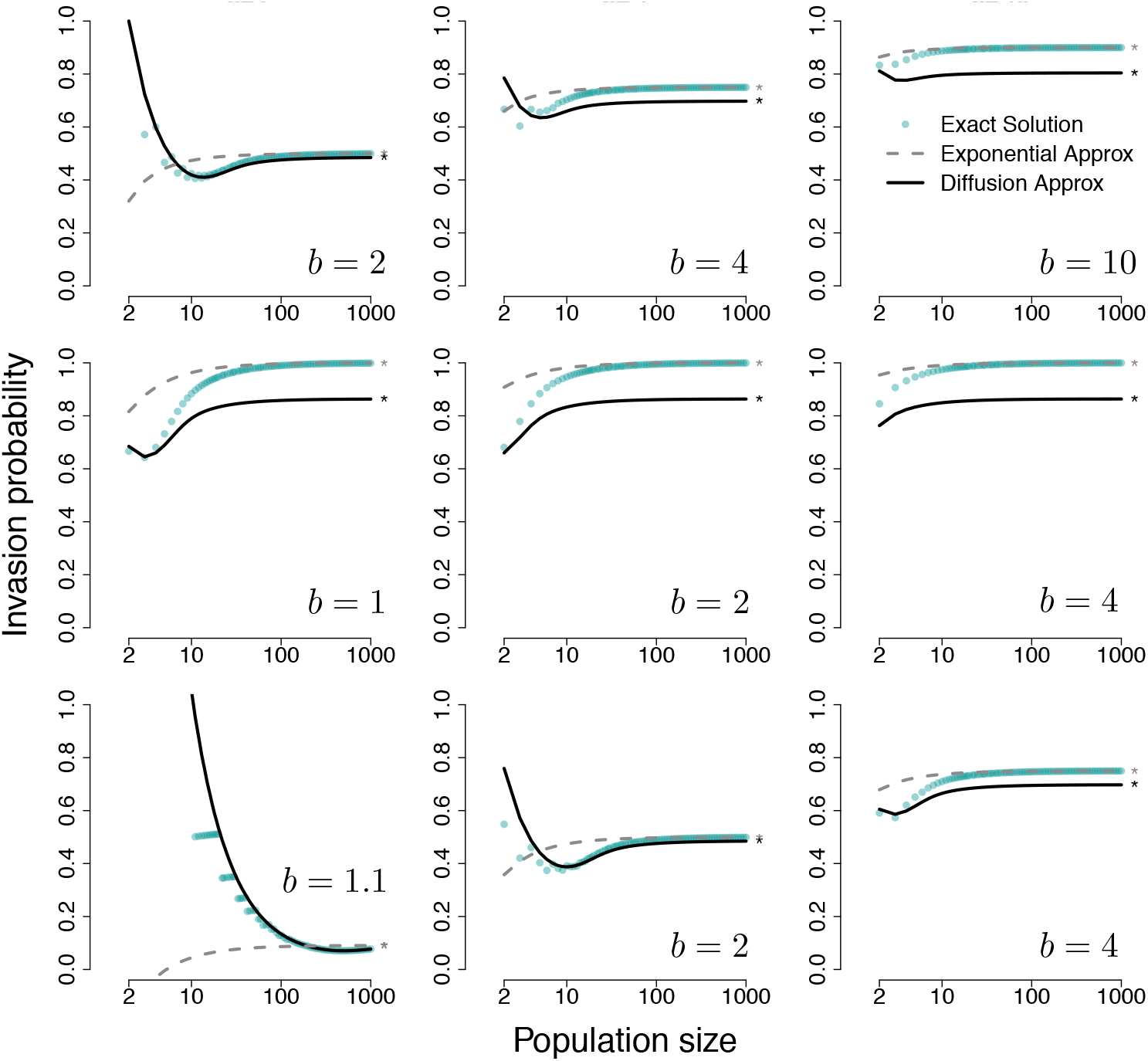
Invasion Probabilities for Examples 1-3. *Comparison of the approximations with the exact solution for the introductory examples: Example stochastic epidemics (β = δ* = 1, *top row), Example density dependent mortality (β* = 1 < *δ = 2, middle row*), *and Example 3 resource-constrained birth (β = δ* = 1, *bottom row*). *In all panels, we set the death rate parameter d* = 1. *For the resource-constrained birth example, we set a* = 1. *Numerical integration was used to calculate the values for both approximation techniques for Example 3.*

For the supercritical systems displayed in Figure 2 the Diffusion Approximation yields a different answer than the exact solution in the large population size limit. This discrepancy between the Diffusion Approximation and the exact solution confirms our asymptotic analysis and is most apparent when the leading coefficients of the birth and death rates are dissimilar. As displayed in the last column of Figure 2, the Diffusion Approximation fails to match the exact solution when the dynamics are far from critical. For population sizes greater than 100 in Figure 3, the Diffusion Approximation fails to match the exact solution for both near critical and far from critical dynamics. The inability of the Diffusion Approximation to approach one is predicted by our asymptotic result that 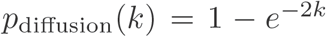 when (*β < δ.* This can be seen in the middle row of Figure 2 and the top row of Figure 3. These plots also highlight the phenomenon that, if the leading order exponents satisfy (*β < δ*, then in the large population limit, the Diffusion Approximation completely ignores the parameters of the rate functions.

By contrast, the Diffusion Approximation succeeds in characterizing the large population size behavior when the dynamics are dominated by the death rate (*µ* is leading order). This is seen in the bottom row of Figure 3, as predicted by our result that 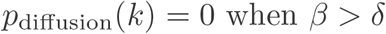 *β < δ*. See Theorem 3.4)

**Fig. 3.**
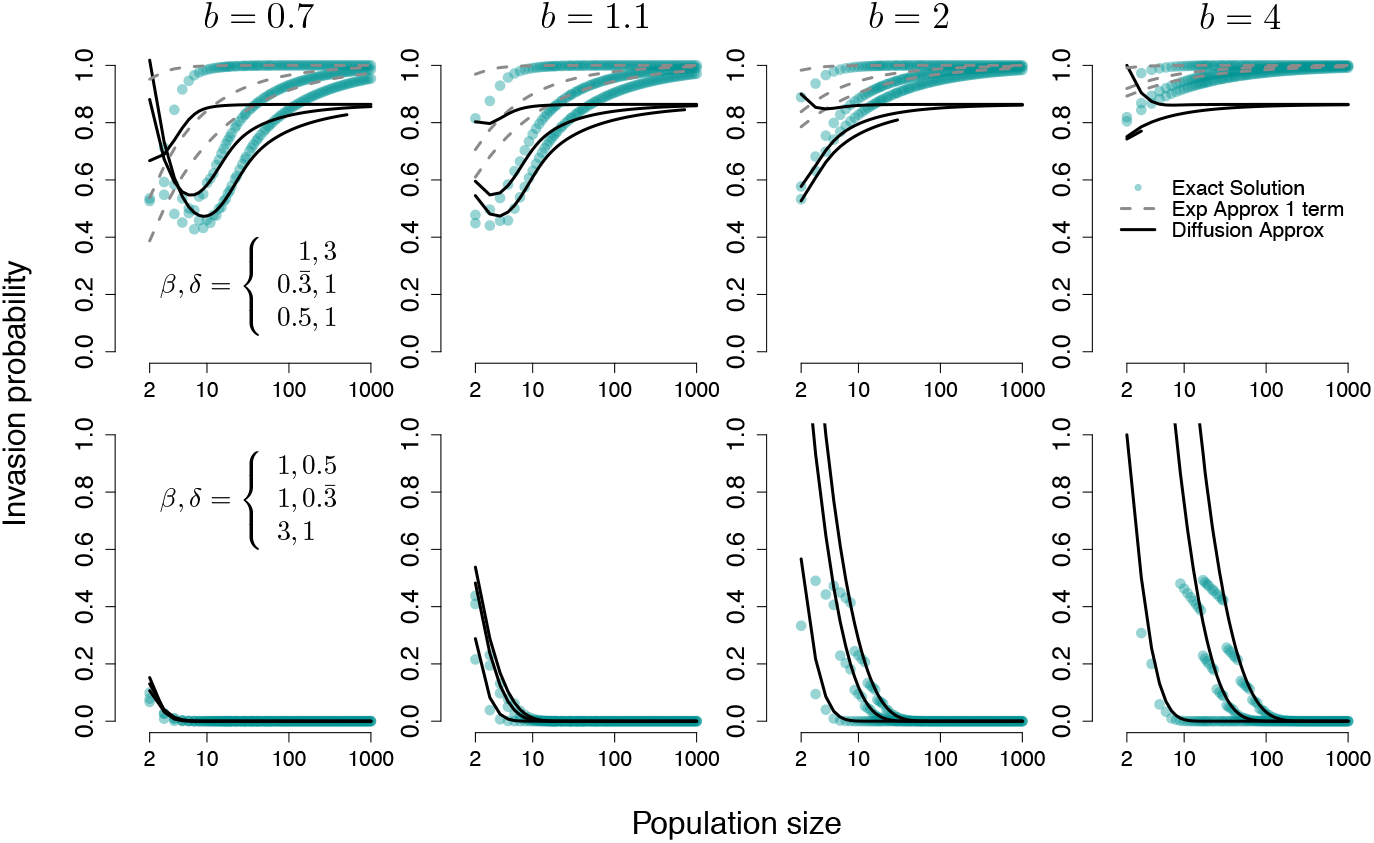
Mismatched Leading Orders. Invasion probabilities for Example 4, in which the birth and death rates are single terms with mismatched exponents. In the top row, the leading order is in the birth rate equation λ(x). In the bottom row, the leading order is in the death rate equation μ(x). Since the Exponential Approximation is invalid in the subcritical case, we have omitted the approximation from the bottom row, where β > δ.

### 5.2. Diffusion approximation methods can work well for small populations that exhibit near critical dynamics

In the examples we have studied, the Diffusion Approximation consistently outperforms the Exponential Approximation when the population is small and the parameter set is near critical. This result is first demonstrated in Figure 2. Moving from left to right in this multi-panel figure, the parameter characterizing the birth rate (b) moves farther away from the death rate parameter (*d*) which is set to one by default. As such, the first column shows that the Diffusion Approximation closely approximates the exact solution for the invasion probability for small population sizes.

The Diffusion Approximation’s transition from exceptional to poor performance is even more clearly demonstrated when studying Example 4. In the top row of Figure 3 we see that the Diffusion Approximation captures non-monotonic features of the exact solution that the Exponential Approximation misses entirely. In this example, the dynamics are dictated by the birth rate since *λ* features the lower leading order term. When the dynamics are near critical *b* = 0.7 and *b* = 1.1, there is a range of small population sizes where the Diffusion Approximation tracks the exact solution.

### 5.3. When leading order terms match, higher order terms matter: for small, but not large population sizes

When the leading order of the “birth” and “death” rates are the same and their leading coefficients are equal, subtleties in the outcomes are determined by the first pair of mismatched coefficients. This result is displayed prominently in Figure 4 for which we chose both rate functions to be asymptotically linear (*β = δ* = 1), to share the same leading coefficient (*b* = *d* = 1), but to have different values for the coefficients *b*_1_ and *d*_1_ (see Assumption 1.3).

**Fig. 4.**
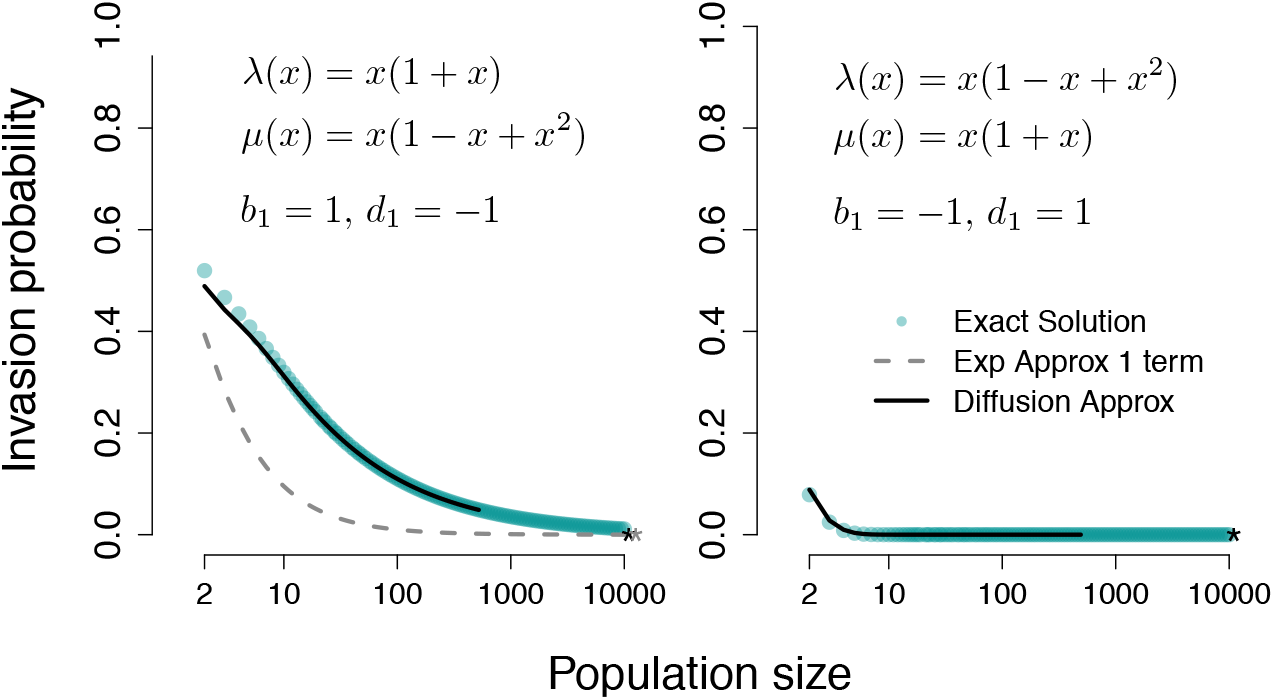
Identical Leading Order Terms. Comparison between the diffusion and exponential approximations when both rate equations are asymptotically linear (β = δ = 1) and the leading coefficients match (b = d = 1). In both panels, the invasion probability tends to zero as the population size (N) becomes large. Differences between the left (supercritical) and right (subcritical) panels are driven by the leading pair of terms with mismatched coefficients (b_1_ ≠ d_1_).

In the left panel of Figure 4, *b*_1_ > *d*_1_ and the system is supercritical (as defined in Assumption 1.1). In this case, both the Diffusion Approximation and Exponential Approximation are well defined. As the population size becomes large, the approximations correctly predict that the invasion probability approaches zero. The corresponding analytical results are presented in Theorems 3.4 and 4.1, respectively, with their asymptotic limits indicated in the plot as black and gray “*”s. Limitations of the numerical integration procedure prevent us from displaying the Diffusion Approximation for the full range of population sizes (i.e. up to 10,000). The main difference between the two approximations in this regime is the rate of convergence to zero, with respect to increases in population size. Consistent with the Diffusion Approximation’s success for relatively small population sizes, it initially tracks the exact solution.

In the right panel of Figure 4, b_1_ < *d*_1_ and the dynamics are subcritical. In Theorem 4.1, we noted that the Exponential Approximation does not hold when *b < d.* As a direct consequence of equation (32), we further observe that the Exponential Approximation will be invalid for sufficiently small population sizes. In particular, when *d = b* and *β = δ*, we have

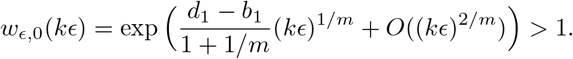

Since the Exponential Approximation is 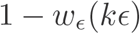, this returns an invalid value less than zero. From a practical point-of-view, one could simply define the Exponential Approximation to be zero in such a circumstance.

### 5.4. Approximation success depends on the initial number of individuals introduced in the population

When more than one individual is initially introduced in a population, the probability of invasion increases. In Figure 5, we display results for each approximation method along with the exact solution for the probability of extinction when 1 ≤ *k* < *Nx*_*_ individuals are initially introduced in the population. By definition, for all larger values of *k*, the invasion probability is one.

**Fig. 5.**
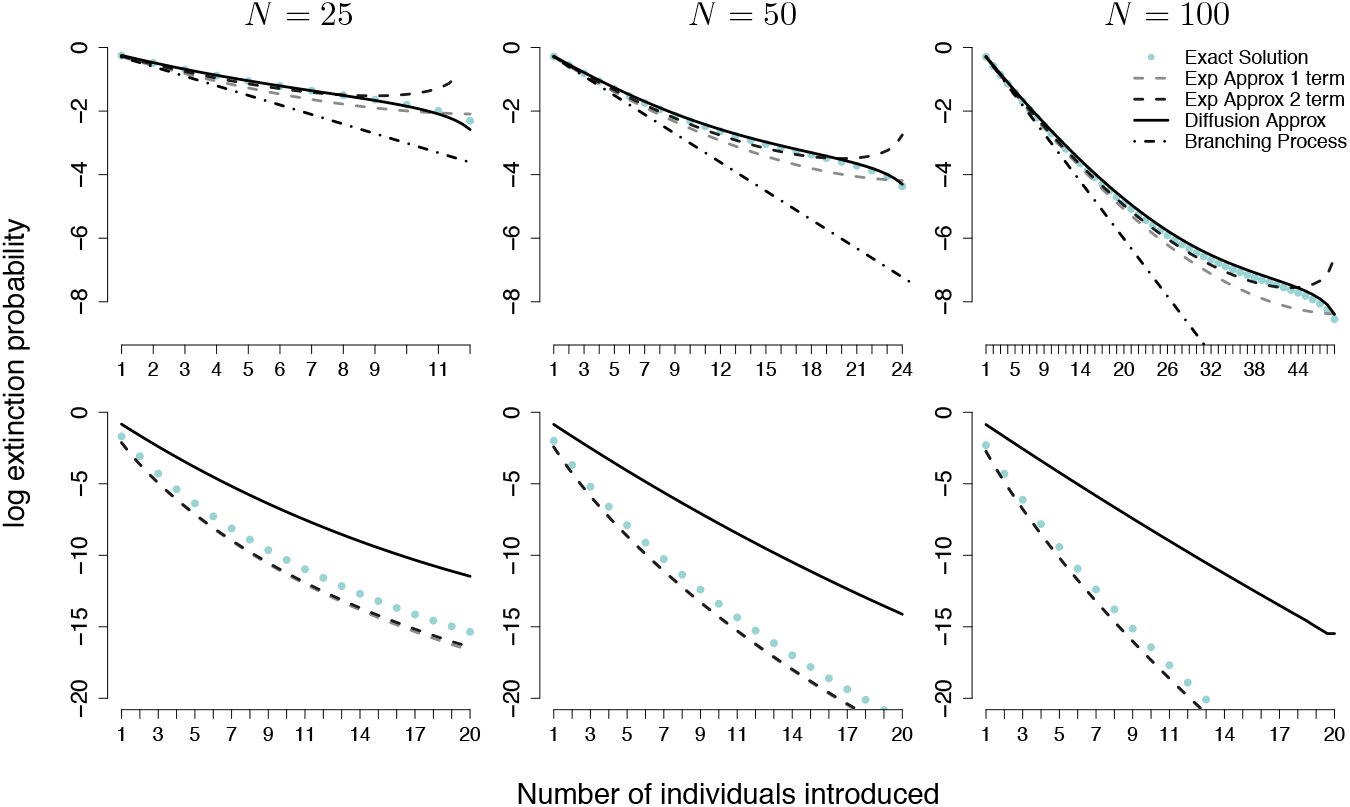
Comparison of the approximations for different numbers of initially introduced individuals. Results are shown for Example 1 (top row) and Example 2 (bottom row) with rate parameters b = 2 and d = 1.

As shown in the top row of Figure 5, when the leading order terms match, the Diffusion Approximation performs well and tracks the exact solution over the full range of initial numbers of individuals introduced. Typically the Exponential Approximation with two terms (dark gray dashed line) better approximates the exact solution than the Exponential Approximation (light gray dashed line). However, when *k* is close to *Nx*_*_ the Exponential Approximation with two terms sharply turns up, away from the exact solution. This numerical result is in line with expectations from our analytical results in equation (28) since the additional higher order term is undefined for *ρ*(*x*) = 1, i.e. when *λ*(*x*) = *µ*(*x*).

In special cases, it is possible to compute the Exponential Approximation by hand using Definition 1.7. We validated the Exponential Approximation for the epidemic extinction probability (Example 1) by comparing the exact result for the leading order approximation,

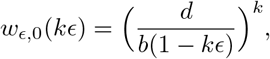

with the numerical integration result plotted in Figure 5. There was not a visible difference between the value found by hand and the numerically obtained value.

We also compared the exact and numerical values for the Exponential Approximation with the analytical approximation found in Lemma 4.3. During this investigation, we found that since the range of potential number of introduced individuals scales with the total population size, it is important to keep track of the *k* parameter in the error term in equation (32). As *k* approaches *Nx*_*_ the error term becomes significant (even for large population sizes). For fixed *k*, the population size can be chosen large enough to yield an approximation with the desired level of accuracy.

## 6. Conclusion

Applied mathematicians, physicists, and biologists use a variety of techniques to approximate Markov chain models for population processes. The widespread use of SDEs, in particular, raises the question of whether these approximations faithfully reproduce results for basic probability questions, especially, hitting times and splitting probabilities. In this work we focused on studying the probability that the lineage of a few newly introduced individuals will invade a larger population. We analyzed the Diffusion Approximation and a simple Exponential Approximation rigorously in the large population limit and numerically for finite size populations. We have were able to show that both population size and the state of sub- versus super-criticality play an important role in determining which approximation method performs better.

Similar to recent analogous results for mean hitting times, we found that, in the large population limit, the Diffusion Approximation does not agree with the asymptotic invasion probability of the original Markov chain system (Theorem 3.4). In fact, when the leading order terms are mismatched (see the case when *β* ≠ *δ* in Theorem 3.4 and Example 2 in the bottom row of Figure 5), the Diffusion Approximation takes on a value that does not depend on the parameters of the Markov chain’s rate functions at all. Interestingly though, when the dynamics are near critical and the population of interest is small, we found that the Diffusion Approximation often performs quite well. This can be seen throughout the figures generated by our numerical investigation, which is described in Section 5. By contrast, for supercritical systems that are far from critical, the Exponential Approximation nearly matches the exact solution for large populations, while the Diffusion Approximation visibly misses the mark. The rigorous expression of this observation can be found in Theorem 4.1. There, we show that for asymptotically linear supercritical systems the Exponential Approximation provides the correct limiting result. This is displayed for a stochastic epidemic model (Example 1) and a resource-constrained population model (Example 3) in Figures 2 and 5.

Our work highlights that invasion probabilities are an important testing ground for evaluating approximation methods. Unlike mean hitting time calculations, the simplicity of the fundamental difference equations determining invasion probabilities permits the explicit evaluation of limits. At this stage, it remains an open question to completely understand which approximation is best for a given set of circumstances. It is possible, but not at all clear, that taking higher order approximations of the Exponential Approximation or a fully formulated WKB-type approximation would succeed under all conditions. In any case, it remains true that careful consideration should be taken when choosing an approximation method for evaluating properties of continuous-time, discrete state-space Markov chains.

## Acknowledgements

The authors would like to thank José M. Ponciano and Sergei S. Pilyugin for thoughtful conversations in the development of this work.

## Appendix A. Series Formulas

*Proposition* A.1 (Series Expansion I). For a given power series with *γ* ≥ 0 and *v* > 0, there exist coefficients 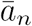 such that

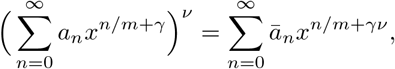

where the first few coefficients are

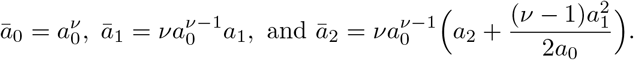

*Proof*. The proof begins by factoring out the leading term in the given series

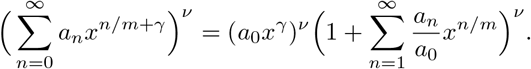

Then make the substitution 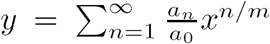 and find the Taylor expansion of 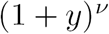,

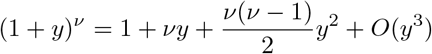

They by plugging in the series for *y* and collecting terms of the same power, we have

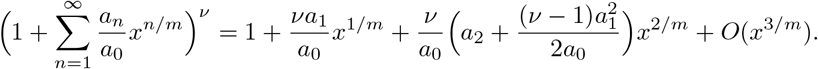

The form of the new series is found by multiplying each term by the original leading order term 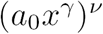.

*Proposition* A.2 (Series Expansion II). Suppose a power series is given in the form, 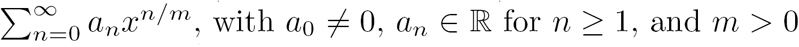. Whenever *x* is such that 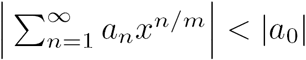, then

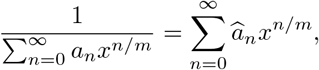

where

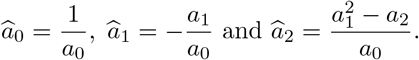

*Proof*. Suppose there is a given power series that satisfies the conditions of the proposition. Further suppose that the inequality 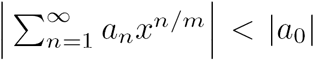 holds. By rearranging our original expression, we find a familiar form

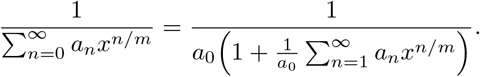

Since we have assumed that 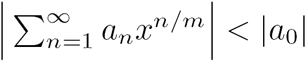,

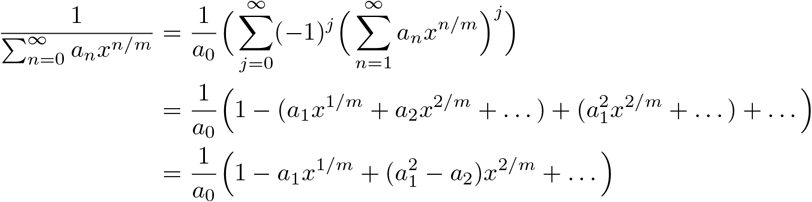

The first few coefficients of the resulting series are obtained by distributing the 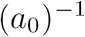 in the last line.

*Proposition* A.3 (Series Inversion [28]). Suppose *f*(*x*) is increasing on [0, *x*_*_) Further assume that *f*(0) = 0 and *f*(*x*) can be written as a formal power series:

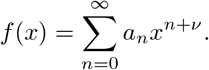

Let *t* = *f*(*x*). Then the inverse of the series *f* can be expressed as

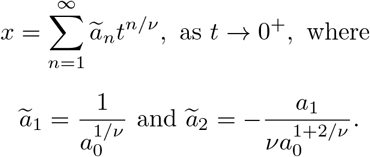

*Proof*. We first rewrite *f*(*x*) as follows

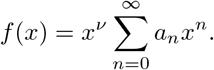

Setting *t* = *f*(*x*), we have

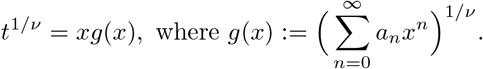

Then, by Proposition A.1, we have

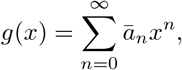

where

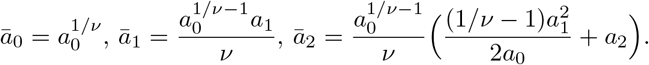

Then,

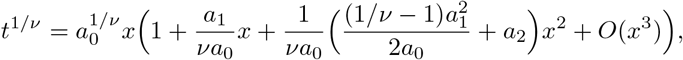

and

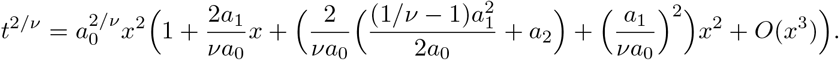

Choose 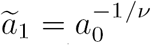 so that it cancels the leading coefficient of 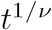. From here we use powers of 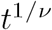 to find an expansion of the form

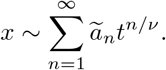

By collecting ordered terms, we find the corresponding two term approximation,

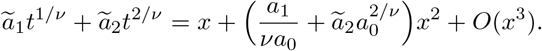

We then choose

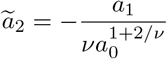

so that the coefficient for the *x*^2^ term is zero. Increasingly precise approximations of *x* can be found by keeping track of the higher order *t* terms and choosing each 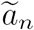 to appropriately cancel out these terms.

